# Brian2GeNN: a system for accelerating a large variety of spiking neural networks with graphics hardware

**DOI:** 10.1101/448050

**Authors:** Marcel Stimberg, Dan F. M. Goodman, Thomas Nowotny

## Abstract

“Brian” is a popular Python-based simulator for spiking neural networks, commonly used in computational neuroscience. GeNN is a C++-based meta-compiler for accelerating spiking neural network simulations using consumer or high performance grade graphics processing units (GPUs). Here we introduce a new software package, Brian2GeNN, that connects the two systems so that users can make use of GeNN GPU acceleration when developing their models in Brian, without requiring any technical knowledge about GPUs, C++ or GeNN. The new Brian2GeNN software uses a pipeline of code generation to translate Brian scripts into C++ code that can be used as input to GeNN, and subsequently can be run on suitable NVIDIA GPU accelerators. From the user’s perspective, the entire pipeline is invoked by adding two simple lines to their Brian scripts. We have shown that using Brian2GeNN, typical models can run tens to hundreds of times faster than on CPU.

## Introduction

GPU acceleration emerged when creative academics discovered that modern graphics processing units (GPUs) could be used to execute general purpose algorithms, e.g. for neural network simulations (Oh and Jung, 2004; Rolfes, 2004). The real revolution occurred when NVIDIA corporation embraced the idea of GPUs as general purpose computing accelerators and developed the CUDA application programming interface (NVIDIA^®^ Corporation, 2018) in 2006. Since then, GPU acceleration has become a major factor in high performance computing and has fueled much of the recent renaissance in artificial intelligence. One of the remaining challenges when using GPU acceleration is the high degree of insight into GPU computing architecture and careful optimizations needed in order to achieve good acceleration, in spite of the abstractions that CUDA offers. Since 2010 we have been developing the GPU enhanced neuronal networks (GeNN) framework (Yavuz et al., 2016) that uses code generation techniques to simplify the efficient use of GPU accelerators for the simulation of spiking neuronal networks.

Brian is a general purpose simulator for spiking neural networks written in Python, with the aim of simplifying the process of developing models (Goodman and Brette, 2008, 2009). Version 2 of Brian (Stimberg et al., 2014) introduced a code generation framework (Goodman, 2010) to allow for higher performance than was possible in pure Python. The design separates the Brian front-end (written in Python) from the back-end computational engine (multiple possibilities in different languages, including C++), and allows for the development of third party packages to add new back-ends.

Here, we introduce the Brian2GeNN software interface we have developed to allow running Brian models on a GPU via GeNN. We analysed the performance for some typical models and find that – depending on the CPU and GPU used – performance can be tens to hundreds of times faster.

## Results

We benchmarked Brian2GeNN on two model networks we will name “COBAHH” and “Mbody” in the remainder of this paper. COBAHH is an implementation of a benchmark network described in Brette et al. (2007). Essentially, this benchmark model consists of *N* Hodgkin-Huxley-type neurons, modified from the model in Traub and Miles (1991), 80% of which form excitatory synapses and 20% inhibitory synapses. All neurons were connected to all other neurons randomly with a connection probability chosen such that each neuron received on average 1000 connections for large models, or connections from all other neurons if the number of neurons was less than 1000.

Mbody is an implementation of the mushroom body model of Nowotny et al. (2005) but unlike in the original publication also with a similar neuron model to the one used for the COBAHH benchmark. The model was used with 100 input neurons, 100 output neurons and varying numbers *N* of Kenyon cells (hidden layer). Projection neurons in the input layer are connected with fixed probability of 15% to Kenyon cells. Up to *N* = 10, 000 Kenyon cells they are connected all-to-all to the output neurons, and for *N* > 10, 000, they are connected randomly with probability chosen such that the output neurons receive input from on average 10, 000 Kenyon cells.

Both models were integrated with an exponential Euler algorithm at 0.1ms time steps. The benchmarks presented here were obtained using the GeNN sparse matrix representation for synaptic connections.

We benchmarked the models on different systems and with different backends. The GeNN backend through the Brian2GeNN interface presented here was compared to the “C++ standalone” backend included with the Brian simulator which runs on the CPU with either a single thread or with multiple threads via the OpenMP interface. Benchmarks were performed for both, single precision (32 bit) and double precision (64 bit) floating point. This is particularly relevant for GPUs because different GPU models have a different number of 64 bit cores, which in addition may be run at reduced clock frequencies for thermal management, and, therefore, can be between only 2× but up to 32× slower in double precision simulations than in single precision (see table 2).

We recorded the overall wall clock time for the simulation including all stages from code generation and initialization in Python to C++ compilation and execution of the binary (“overall runtime”). We also took more fine-grained measurements of the time for code generation and compilation, the time spent for synapse creation and initialization, the time spent for the actual simulation and the overhead, including, e.g., time spent on reformatting data structures between Brian 2 and GeNN formats, copying to and from the GPU and writing results to disk.

### Simulation time

The results for the net simulation time for the two models on CPU and the TITAN Xp and V100 GPUs are shown in figure 1 as a function of the size of the models, indicated by the total number of neurons. GeNN offers two different strategies for parallelising the spike propagation logic, along pre-synaptic inputs (looping over post-synaptic targets) or along post-synaptic targets, looping over pre-synaptic sources. We benchmarked both algorithms for each of the models.

**Figure 1:**
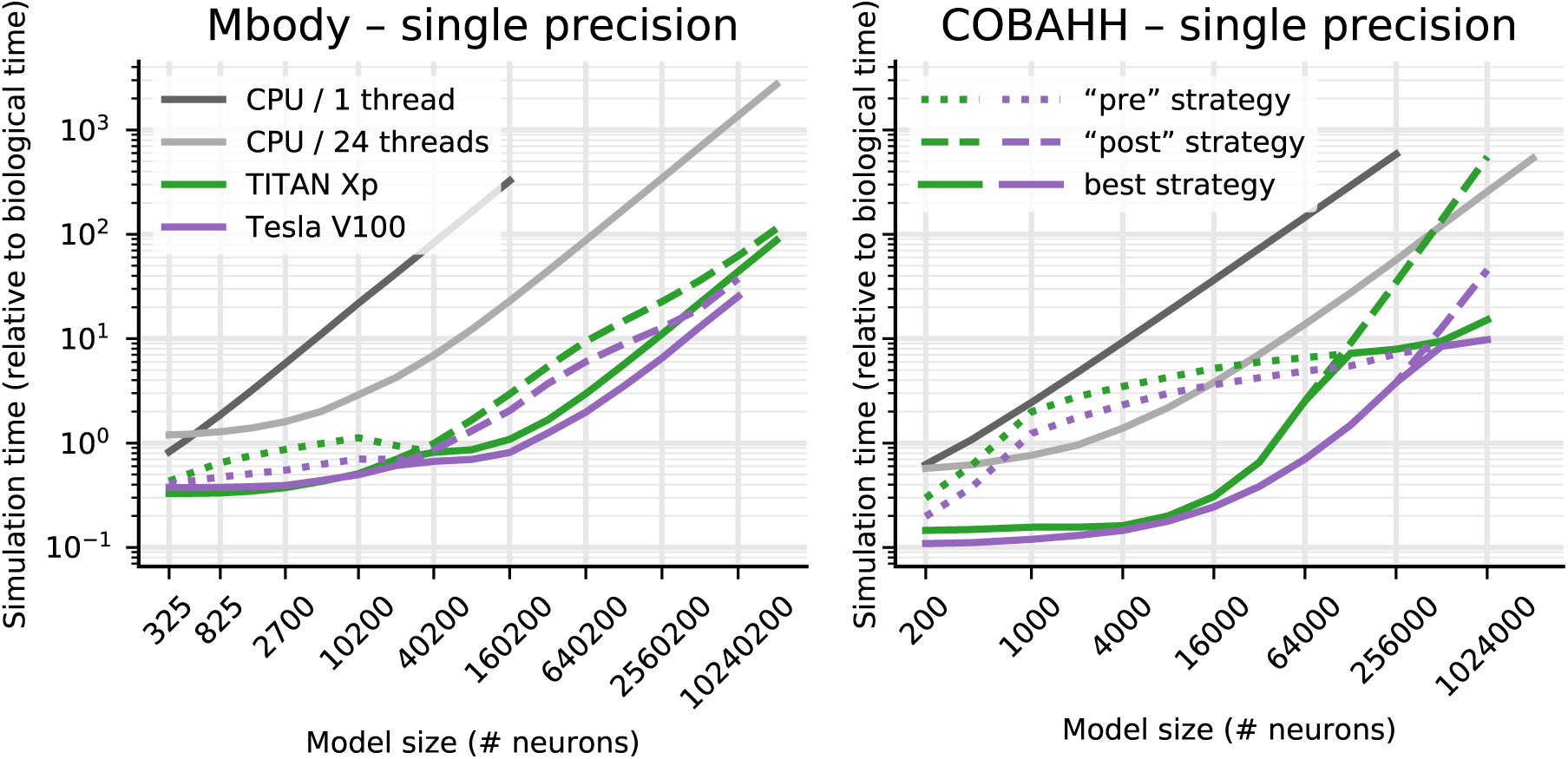
Benchmark of the net simulation time on a 12 core CPU with a single thread (dark gray) or using OpenMP with 24 threads (light gray), compared to a consumer GPU (TITAN Xp) and an HPC model (V100). For the GPUs, simulation times are displayed separately for a pre-synaptic parallelisation strategy (dotted) or post-synaptic strategy (dashed). The better of the two strategies is highlighted by a solid line.

As is often the case, the single thread CPU solution scales essentially linearly with the size of the two models, expressed in terms of the number of simulated neurons (figure 1). This reflects the linear scaling of processing time with the number of operations required and that both models are essentially neuron-bound on the CPU due to their computationally expensive neuron model, their chosen connectivity and the observed number of spikes. The 24-thread OpenMP simulations take initially the same time for very small models but we found that the simulations ran about 13–14 times faster than on a single CPU core for the largest MBody model tested on single CPU core (160,200 neurons) and 8–11 times faster for the largest COBAHH model testes on single CPU core (256,000 neurons). Larger models were only tested on 24-thread OpenMP and GPUs due to the prohibitively long runtime on a single CPU core. For models larger than 40,200 neurons (Mbody) and 8000 neurons (COBAHH), the 24 thread OpenMP solution also scales approximately linearly with the number of neurons.

The simulations run on the GPU via Brian2GeNN (green and purple lines in figure 1) were significantly faster than the 24 thread OpenMP (light gray), for instance, 40–54 times faster in the Mbody model for 10,240,200 neurons and up to 24–26 times faster in the COBAHH model for 1,024,000 neurons when using the V100 GPU. We have summarised the observed speed-ups achieved for the simulation time in table 1. Overall the GPU runs always faster than a single threaded CPU version, up to a factor of 400, but when compared against the 24 thread OpenMP version, acceleration can vary from 2× slower than the CPU to about 50× faster.

**Table 1:** Acceleration over simulation on CPU. This only considers simulation time. Numbers are relative to simulations on the host of the Titan Xp GPU (see table 2) and compare to a single-thread simulation (left) or a 24-thread OpenMP simulation (right). The two numbers shown are for single precision (gray background) and double precision (white background).

**Mbody benchmark**
| # neurons | *compared to CPU 1 thread* |  |  |  |  |  | *compared to CPU 24 thread* |  |  |  |  |  |
| --- | --- | --- | --- | --- | --- | --- | --- | --- | --- | --- | --- | --- |
|  | 40,200 |  | 80,200 |  | 160,200 |  | 40,200 |  | 160,200 |  | 10,240,200 |  |
| Quadro K2200 | 39.2 | **6.5** | 52.0 | 6.7 | 60.7 | 7.0 | 3.3 | 0.6 | 4.3 | **0.5** | 4.3 | **0.5** |
| Tesla K40c | 34.7 | 25.6 | 58.8 | 39.8 | 80.9 | 53.8 | 2.9 | 2.4 | 5.7 | 4.2 | 7.1 | 5.8 |
| Titan Xp | 101.9 | 39.4 | 190.3 | 51.4 | 300.3 | 60.9 | 8.5 | 3.7 | 21.1 | 4.8 | 31.0 | 5.0 |
| Tesla V100 | 124.3 | 105.9 | 235.5 | 191.7 | **401.6** | 251.7 | 10.4 | 9.9 | 28.3 | 19.7 | **53.9** | 40.4 |

**COBAHH benchmark**
| # neurons | 64,000 |  | 128,000 |  | 256,000 |  | 64,000 |  | 256,000 |  | 1,024,000 |  |
| --- | --- | --- | --- | --- | --- | --- | --- | --- | --- | --- | --- | --- |
| Quadro K2200 | 18.0 | 4.9 | 24.2 | 4.5 | 13.6 | **4.1** | 1.7 | 0.6 | 1.3 | **0.5** | — | — |
| Tesla K40c | 20.6 | 16.3 | 11.7 | 9.4 | 16.7 | 8.6 | 1.9 | 1.9 | 1.6 | 1.1 | 1.0 | — |
| Titan Xp | 57.7 | 29.1 | 40.3 | 22.2 | 73.8 | 33.7 | 5.4 | 3.4 | 7.2 | 4.1 | 16.9 | 6.5 |
| Tesla V100 | **207.7** | 155.7 | 196.5 | 134.4 | 154.1 | 113.5 | 19.6 | 18.3 | 15.0 | 13.9 | **26.3** | 24.2 |

**Table 2:** Configurations used for benchmarking.

| CPU | # cores | Clock speed (GHz) | Memory (GB) | GPU | Architecture | # cores | Memory (GB) | Performance* (float) | Performance* (double) |
| --- | --- | --- | --- | --- | --- | --- | --- | --- | --- |
| **Intel Xeon E5-1630 v3** | 4 | 3.7–3.8 | 16 | **Quadro K2200** | Maxwell | 640 | 4 | 1,439 | 45 |
| **Intel Xeon E5-1620 v2** | 4 | 3.7–3.9 | 32 | **Tesla K40c** | Kepler | 2,880 | 12 | 4,290 | 1,430 |
| **Intel Core i9-7920X** | 12 | 2.9–4.4 | 64 | **TITAN Xp** | Pascal | 3,840 | 12 | 12,150 | 380 |
| **Dual Intel Xeon Gold 6148** | 2 × 20 | 2.4 | 192 | **Tesla V100** | Volta | 5,120 | 16 | 14,131 | 7,066 |

\*maximum performance in GFLOPS

Interestingly, the different parallelisation methods for spike propagation available in GeNN (dashed and dotted lines in figure 1) perform differently as a function of size and of the two models being simulated. The post-synaptic method is always faster for small models while the pre-synaptic method wins for very large models. However, for the Mbody example, the swap occurs at moderate model sizes of about 40,200 neurons, whereas for the COBAHH model, it is for much larger models (128,000 neurons for the TITAN Xp and 512,000 neurons for the V100). Also, while the differences of the two methods are not that pronounced for the large Mbody models, the post-synaptic method in the COBAHH model scales very poorly with size at large model sizes, leading to quite low performance of Brian2GeNN in this mode. The pre-synaptic method, on the contrary, is not particularly fast for smaller to medium sized COBAHH models (even slower than the 24 thread OpenMP version), but scales excellently for the largest models, leading to significant speedups over OpenMP.

The simulation times for a larger variety of different GPU hardwares are shown in figure 2. Note that we here display the results for the better of the two parallelisation strategies for each model run. We benchmarked four different graphics cards (see table 2). The results show consistent trends given the specifications of the hardware (table 2), even though some may not be as obvious as others. The V100 is almost always fastest, typically followed by the TITAN Xp, K40 and Quadro card in this order. Note however, the marked difference in double precision performance for the consumer cards (Quadro and TITAN Xp), compared to the high performance computing cards (K40c and V100). This is expected because the consumer cards have NVIDIA GPU architectures (Maxwell respectively Pascal) that have fewer double precision cores and double precision operations are hence up to 32 times slower than single precision, while the HPC cards used here are Kepler and Volta architecture and have only a factor 2 performance difference between double precision and single precision operations. Accordingly, while in single precision, the presumably less powerful but more recent Quadro card performs at the level of or even better than the older but larger K40c accelerator, it does not compare favourably for double precision.

**Figure 2:**
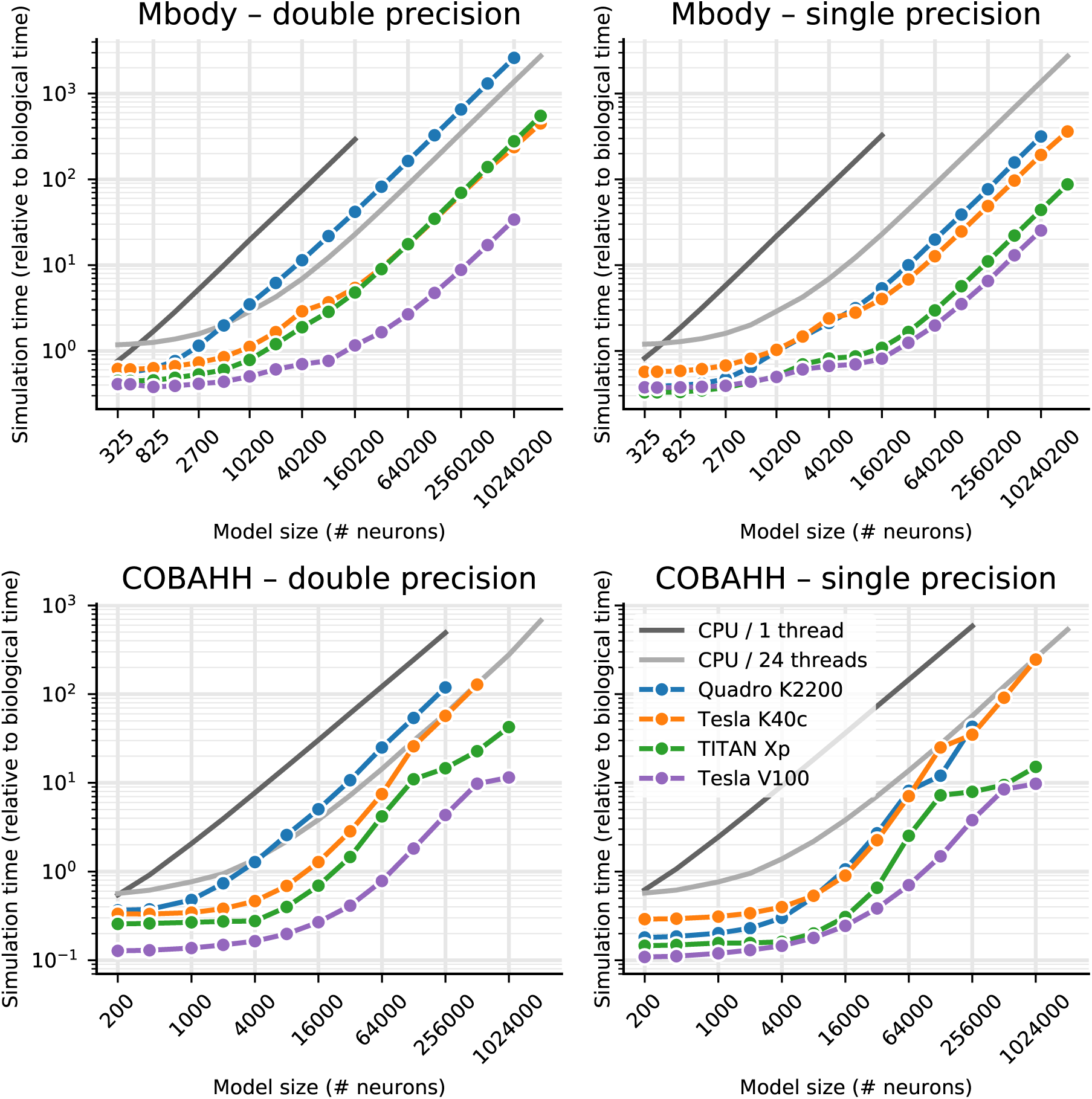
Benchmarking of the net simulation time for different GPU models. Measurements were taken separately for the MBody model (top) and COBAHH model (bottom) for double precision floating point (left) and single precision (right). Simulation time is shown relative to the simulated biological time. CPU performance was measured on the host of the TITAN Xp GPU (see table 2). For the GPUs, the better (smaller) of the simulation times for either pre-synaptic or post-synaptic parallelisation strategy are shown. See figure 1 and main text for more in-depth explanation.

Comparing the two models, it is clear that the performance gains of Brian2GeNN on the different GPU platforms is more marked for the Mbody model than for the COBAHH model. This would be expected for the spike propagation code because the mainly feedforward structure of the Mbody model lends itself better to parallelisation on GPUs with GeNN than the randomly recurrently connected COBAHH model. Inspection of the computation time used for neuron and synapse updates and synapse updates (spike propagation) revealed that for both models, but contrary to the CPU situation, a large percentage of computing time is typically spent for spike propagation when using a GPU.

### Time for other tasks

So far we have presented results for the core simulation time. As explained in the methods, Brian2GeNN has a substantial pipeline of tasks before and after the main simulation takes place. Figure 3 illustrates the essence of how the computation times necessary along this pipeline stack up. We defined four main phases of a Brian2GeNN run: “code generation and compilation”, “synapse creation”, “main simulation” and “overheads”, which bundles smaller tasks such as transforming data formats between Brian 2 format and GeNN format, copying from and to the GPU and writing results to disk. For illustration we have used the data from the TITAN Xp card. The data in the top two panels in figure 3 repeats the results for the simulation time but also shows extrapolations for shorter and longer runs, where computation times are strictly proportional to the length of simulated biological time. The bottom two panels show the compute time spent on the other three phases. Code generation and compilation is a fixed cost that is completely independent of the model size. On the contrary, computation time for synapse creation and initialisation increases linearly with model size in terms of the number of neurons. The other overheads are initially almost independent of model size but then also start increasing with the number of neurons. In the balance, for small to mid-sized models and short simulation runs (1s biological time), code generation and compilation dominates the overall runtime whereas for large models and longer runs, the time for the main simulation dominates.

**Figure 3:**
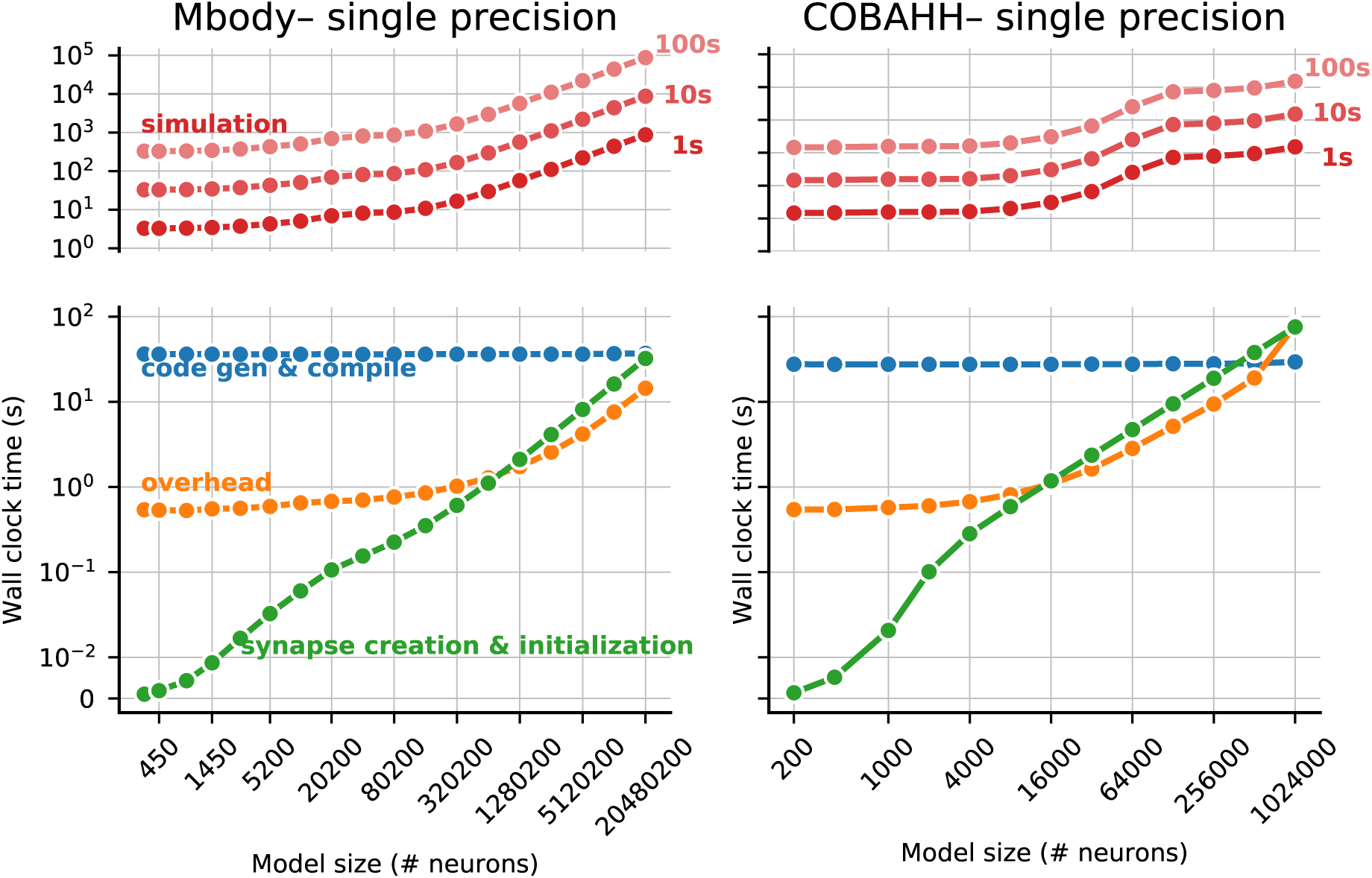
Overview of the components that make up the total runtime of a simulation for the Mbody (left) and the COBAHH benchmark (right). The top panels show the time spent in the simulation itself which scales with the biological runtime of the model (shown at the right) and dominates the overall runtime for big networks and/or long simulations. The bottom panels show the time spent for code generation and compilation (blue), general overhead such as copying data between the CPU and the GPU (orange), and the time for synapse creation and the initialization of state variables before the start of the simulation (green). The details shown here are for single-precision simulations run on the Titan Xp GPU.

To give a rough guide at which amount of biological time for any given model size it becomes viable to use Brian2GeNN we have calculated the minimum simulated biological time for which the overall runtime for Brian2GeNN is smaller than a 24 thread OpenMP solution (figure 4). For simulated biological time of 100s or more it is always faster to use Brian2GeNN, regardless of model size or employed GPU accelerator. For shorter simulated time it depends on the simulated model and the GPU. For example, simulating 10s biological time is perfectly viable on a V100 for the Mbody model at size 40,200 but would be slower on a K40c; or, simulating 10s biological time would not be viable for any of the tested GPUs for the COBAHH model at size 8,000 but viable on all of them at size 64,000.

**Figure 4:**
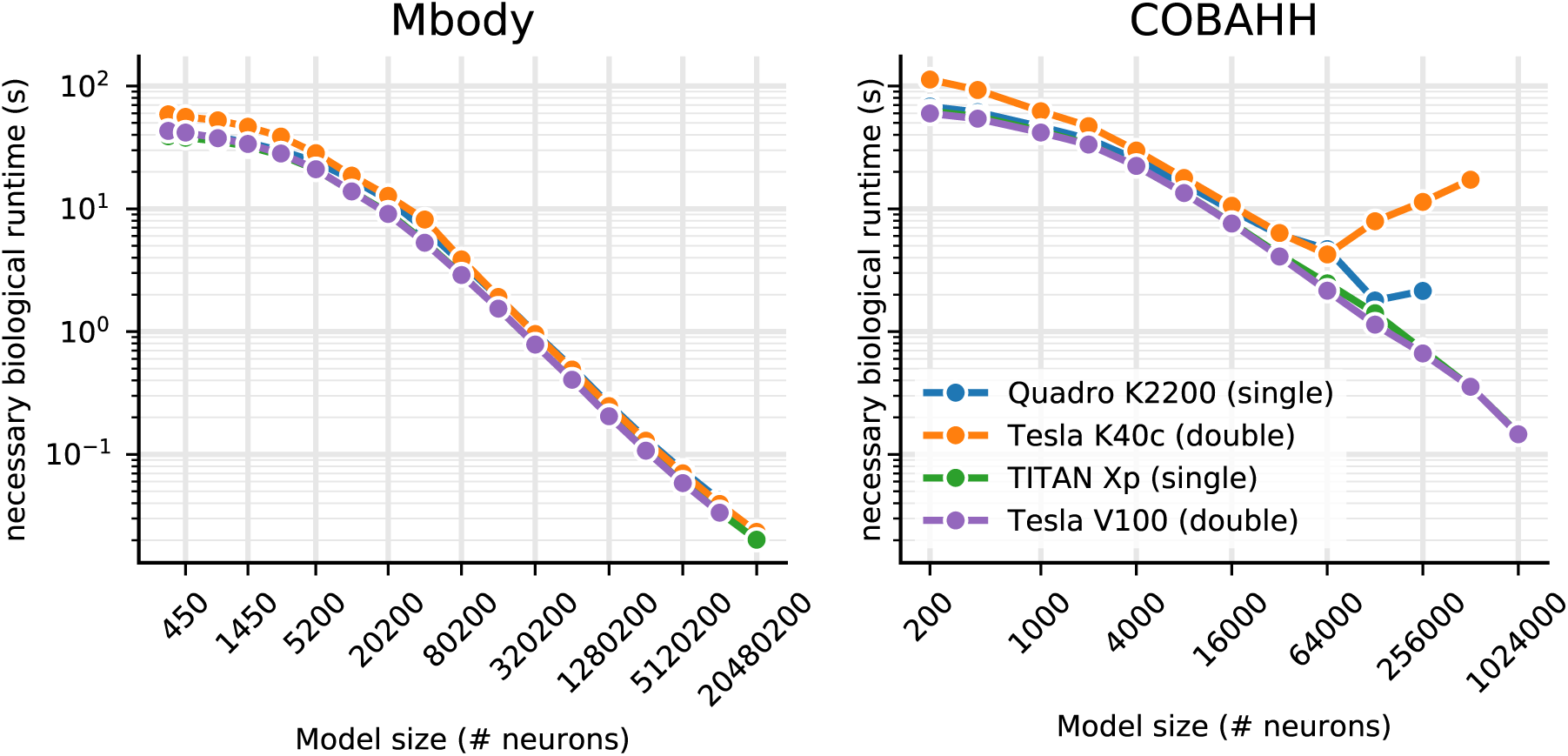
Minimal biological runtime after which the total simulation time, including preparations such as code generation and compilation (cf. figure 3), is smaller when using a GPU compared to 24 threads on a CPU, for networks of different sizes. The CPU comparison is the host of the Titan Xp GPU (see table 2). Results for the Mbody benchmark (left) and the COBAHH benchmark (right). The calculations are based on single precision performance for the Quadro GPU (blue) and Titan Xp GPU (green), and on double precision performance for the Tesla K40c (orange) and the Tesla V100 GPU (purple).

## Discussion

In designing software for computational neuroscience, there are two seemingly conflicting requirements: for high performance and for flexibility. The ability to create new types of models is essential for research that goes beyond what is already known at the time that a simulator package is created. However, hand written code that implements particular models can be much more computationally efficient. This is particularly true in the case of GPU simulations, as it is difficult to make maximally efficient use of GPU resources. Consequently, almost all GPU-based simulators for spiking neural networks have not made it possible to easily create new user-defined neuron models (Nageswaran et al., 2009; Fidjeland and Shanahan, 2010; Mutch et al., 2010; Hoang et al., 2013; Bekolay et al., 2014). The exceptions are GeNN, the package Brian2CUDA (Augustin et al., 2018) currently under development, and ANNarchy (Vitay et al., 2015), which is discussed below.

The technique of code generation allows us to solve this apparent conflict, and has been used by both the GeNN and Brian simulators (Goodman, 2010; Yavuz et al., 2016; Stimberg et al., 2014). In the case of GeNN, when writing a new model users need to write only a very small section of generic C++ code that defines how the variables of a neuron model are updated, and this is then inserted into a detailed template that allows that model to be simulated efficiently on a GPU. Brian meanwhile allows users to write their model definition at an even higher level, as standard mathematical equations in a Python script. These are then automatically converted into low-level C++ code to be compiled and executed on a CPU. In both cases, users write high level code (short snippets of C++ in the case of GeNN, or Python/mathematics in the case of Brian) and efficient low level code is automatically generated.

Linking Brian and GeNN accomplishes two tasks. Firstly, it allows existing Brian users to make use of a GPU to run their simulations without any technical knowledge of GPUs (via GeNN). Secondly, it gives GeNN users a high level and feature packed interface (Brian and Python) to manage their simulations. GeNN was originally designed to be used at the C++ level, with network setup and simulation management handled by the user in C++, but not all computational neuroscientists are comfortable working at this level and there can be considerable savings in development time working at a higher level.

### Related work

The only other spiking neural network simulation package to allow for flexible model definition in a high level language, and for code to run on GPUs, is ANNarchy (Vitay et al., 2015). This simulator was originally designed to adapt a model definition syntax similar to Brian’s to rate-coded networks (rather than spiking neural networks), and to make use of GPUs for high performance. It has subsequently been updated to allow for the definition of spiking neural networks as well as hybrid networks, and simulating spiking networks on the GPU is now provided as an experimental feature. In contrast to Brian2GeNN which supports all major operating systems, ANNarchy only supports running simulations on the GPU on Linux.

As noted in Brette and Goodman (2012), on GPUs it is unlikely that there is a single best algorithm for spiking neural network simulation, but rather the best algorithm will depend on the model. A diversity of GPU spiking neural network simulator packages is therefore desirable.

### Limitations

Brian’s framework for defining models of neurons, synapses, networks and computational experiments is designed to be as expressive and flexible as possible. Consequently, not all features of Brian are available in GeNN, and not all simulations that can be run in GeNN will run efficiently. Among the most important currently unsupported features are continuous, i.e. not spike-based, connections (used for example to implement electrical synapses); heterogeneous, i.e. synapse-specific, synaptic delays; arbitrary, time-varying continuous stimuli; and complex simulation schedules (for example, multiple simulation runs or different simulation time steps for individual groups of neurons/synapses). Attempting to use an unsupported Brian feature with Brian2GeNN will simply raise an error.

However, some features that are supported may also lead to slow code on the GPU. This is because efficient use of the GPU requires appropriate paralellisation strategies and specific memory access patterns, and for some features (particularly relating to backpropagation of information in synapses) it is very difficult to arrange data in memory so that it can be accessed efficiently for both, forward and backward propagation on the GPU (Brette and Goodman, 2012). The very different scaling of runtimes in the COBA example for pre- and post-synaptic parallelisation strategies for synaptic updates in large model instances, as seen in figure 1, is a very typical example of such phenomena. However, it is not straightforward to predict when problems of this kind will be significant. The Mbody example has STDP but because it is otherwise well suited for GeNN due to essentially feedforward connectivity for the majority of synapses and sparse firing, it speeds up well in Brian2GeNN. The COBA example does not have plasticity and yet, due to its relatively dense, random connectivity and somewhat higher firing rates, the speedups are good but less pronounced than in the Mbody example.

### Future work

Further work on Brian and GeNN will go in two main directions. On the GeNN side, we plan to expand the features available in GeNN to cover more of the features available in Brian, as well as improving efficiency. A specific bottleneck that has been recently identified is the synapse creation task (see figure 3). Work is under way that enables synapse creation on the GPU instead of the CPU with considerable performance advantages, in particular where synaptic connectivity becomes more intricate.

On the Brian side, we plan to simplify and optimise the process of writing third party back-ends. This will not only simplify future development of Brian2GeNN but will also encourage the development of an ecosystem of back-ends, for example featuring different GPU algorithms or targeting different computational hardware such as field programmable gate arrays (FPGAs). An interface to generate CUDA code directly from a Brian script, called Brian2CUDA (Augustin et al., 2018), is also under development, but has not yet been released.

For Brian2GeNN itself, we are planning to expose more of the optimisation choices offered to direct users of GeNN to Brian2GeNN users, for instance per-synapse group choices for connectivity matrix representations (sparse, dense, ragged, bitmask) and parallelisation strategies (pre- or post-synaptic). We will also work on exposing the emerging on-GPU initialisation methods mentioned above and the heterogeneous synaptic delays that were recently introduced to GeNN.

## Methods

### Brian2GeNN

Brian2GeNN makes use of the existing code generation facilities in the Brian and GeNN simulators. These code generation facilities differ in important aspects. The Brian simulator provides a comprehensive code generation framework that converts not only high-level descriptions of neural and synaptic models to executable code, but also extends this framework to model initialization including the generation of synapses according to high-level rules. In addition, the user code is written in Python, a language that is very accessible to researchers with a less technical background. However, the generated code is C++ code that runs only on the CPU, and therefore cannot make use of the computational power of GPU accelerators. GeNN’s code generation framework on the other hand is focused more on organizing the code to run efficiently on highly parallel GPUs, leaving the task of defining the code for simulating the neural and synaptic model, and the details of how to run the overall simulation to the user. This is completed in C++, which allows tight integration with other C++ based code, e.g. in the context of robotic controllers, but also makes writing a GeNN simulation relatively difficult for inexperienced programmers. The major advantage of using GeNN is its ability to generate efficient CUDA code that can be executed on a GPU to accelerate simulations.

Brian2GeNN acts as a “glue” between Brian and GeNN, thereby combining the advantages of both simulators. It is built as an extension of Brian’s code generation mechanism and can therefore be directly used from within a Brian script; by choosing the “GeNN device” (l. 2–3, figure 5 top), a standard Brian simulation is turned into a hybrid Brian/GeNN simulation. Such a script typically sets up the simulation components and then triggers the simulation of the network (figure 5 top and bottom left). At this point, the code generation process is activated and generates, compiles and executes the target code. The results of this simulation are then written to disk by the executed code, enabling the Python code to access the requested results to analyze or plot them. The executable code (figure 5 bottom right) is jointly generated by Brian (blue boxes), Brian2GeNN (green boxes/arrows), and GeNN (red box) and executed partly on the CPU and partly on the GPU. The initial steps, synapse creation and model initialization, are unchanged from Brian’s default code generation process. However, since Brian and GeNN use different data structures to represent synapses, Brian2GeNN has to generate code to convert between the two formats. In addition, it copies all the data to the GPU so that it can be used during the simulation run. The main simulation loop delegates the core of the simulation, the dynamic update of neuronal and synaptic state variables as well as the propagation of synaptic events, to the code generated by the GeNN simulator, which executes on the GPU. After each time step, some of this data may be copied back from the GPU and converted to the Brian format so that it can be recorded by Brian’s monitoring mechanism. After the end of the simulation run, Brian2GeNN takes care to copy all data back from the GPU and to convert it to the Brian format, so that Brian can store the results to disk and make them available for analysis in the Python script.

**Figure 5:**
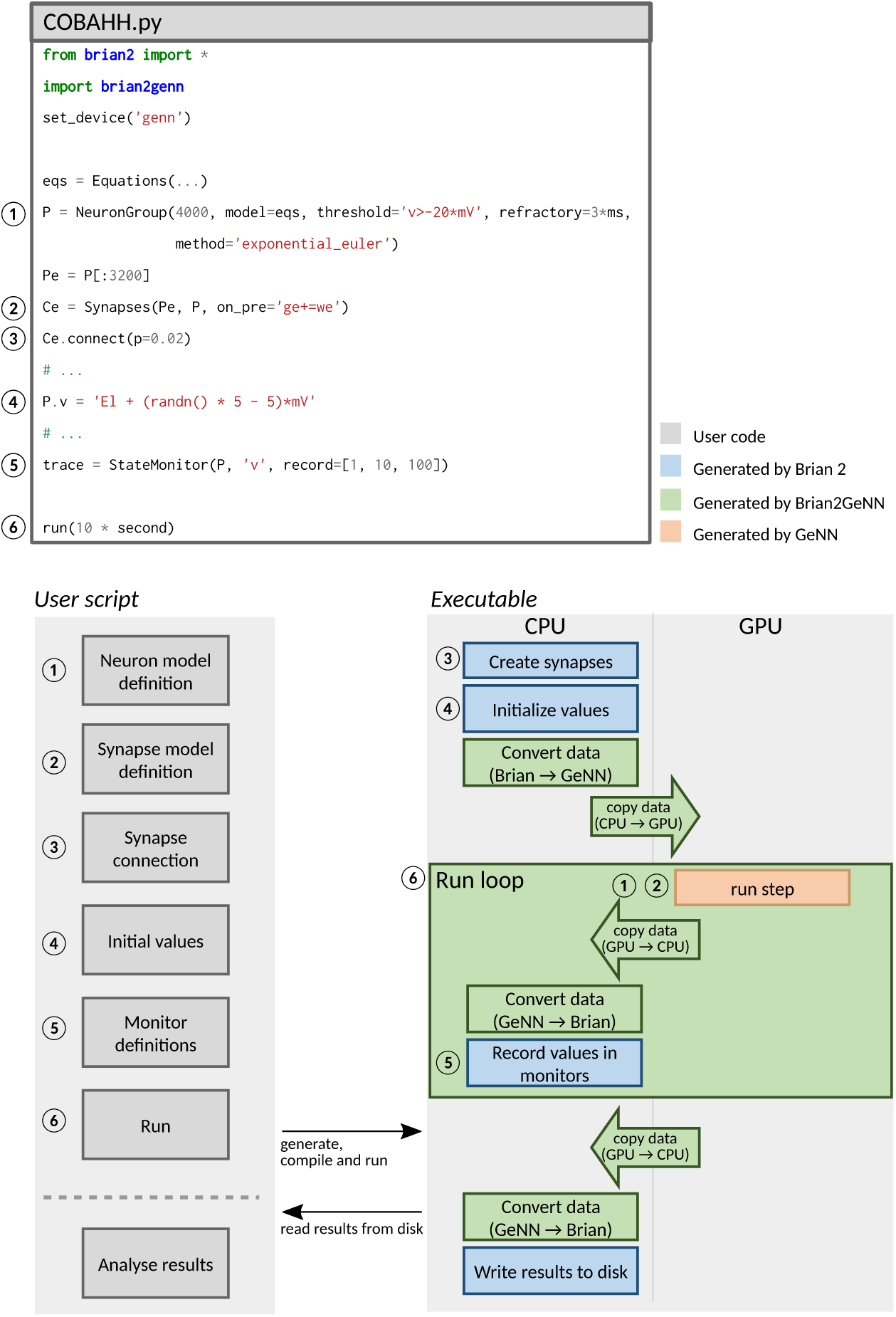
Running simulations with Brian2GeNN. **Top:** Excerpt from an example Brian script that will execute in a hybrid Brian/GeNN simulation due to the import of the `brian2genn` library (line 2) and the selection of the “GeNN device” (line 3). **Bottom left:** General workflow of a Brian2GeNN simulation: the run call triggers the code generation, compilation and execution. After the successful run, results are stored to disk and made available to the Python script. **Bottom right:** Structure of generated code. Parts of the code result from Brian’s standard code generation process (blue), while the main run step is implemented by GeNN (red) and everything is connected together by Brian2GeNN (green). The preparation of the simulation and actions such as variable monitoring are executed on the CPU (left), while the core of the simulation is executed on the GPU (right).

### Benchmarks

Benchmarks were run on a number of different workstations, with different GPUs installed ranging from a standard consumer card (Quadro K2200) to a more powerful gaming GPU (TITAN Xp), an older computing accelerator model (Tesla K40c) to the most recent and most powerful accelerator (Tesla V100). The different configurations for benchmarking are listed in table 2. We used Brian 2, version 2.2 (Stimberg et al., 2018a), GeNN version 3.2 (Knight et al., 2018), and Brian2GeNN version 1.2 (Stimberg et al., 2018b) for our benchmarks.

All benchmarks were run without “monitors”, Brian’s mechanism for recording the activity during a simulation, as we observed in exploratory tests that monitors play only a minor role in the context of the two models used as benchmarks here. For the runs using Brian2GeNN, we used GeNN’s Yale sparse matrix representation (Yavuz et al., 2016) throughout. While for smaller models, dense matrix representations may have speed advantages, the more relevant mid- and large-scale models would lead to “out of memory” failure on all tested GPUs with either of GeNN’s dense matrix representations. Even with sparse matrix representation, some runs failed because of memory overrun. The corresponding data points were omitted from the benchmark figures below.

## Acknowledgments

We thank James Knight for assisting us with running benchmarks on the V100 device and helping with adjustments in GeNN. This work was partially funded by the EPSRC (grants EP/J019690/1, EP/P006094/1) and Horizon 2020 research and innovation program under grant agreement n° 85907 (Human Brain Project, SGA2).

## Author contributions statement

Author contributions: MS, DG and TN developed Brian2Genn, MS, DG and TN ran benchmarks, MS produced figures, MS, DG and TN wrote the manuscript. All authors reviewed the manuscript.

## Additional information

Brian2GeNN is developed publicly on github (https://github.com/brian-team/brian2genn). The scripts and raw results of the benchmark runs are available at https://github.com/brian-team/brian2genn_benchmarks.

The authors declare that they have no competing interests with respect to this work and the funders have not played any role in its design or interpretation.

## References

Augustin, M., Alevi, D., Stimberg, M., and Obermayer, K. (2018). Flexible simulation of neuronal network models on graphics processing units: an efficient code generation approach based on brian. In Bernstein Conference 2018.

Bekolay, T., Bergstra, J., Hunsberger, E., DeWolf, T., Stewart, T. C., Rasmussen, D., Choo, X., Voelker, A. R., and Eliasmith, C. (2014). Nengo: a Python tool for building large-scale functional brain models. Frontiers in Neuroinformatics.

Brette, R. and Goodman, D. F. M. (2012). Simulating spiking neural networks on GPU. Network (Bristol, England), 23(4): 167–82.

Brette, R., Rudolph, M., Carnevale, T., Hines, M., Beeman, D., Bower, J. M., Diesmann, M., Morrison, A., Goodman, P. H., Harris, F. C., Zirpe, M., Natschläger, T., Pecevski, D., Ermentrout, B., Djurfeldt, M., Lansner, A., Rochel, O., Vieville, T., Muller, E., Davison, A. P., El Boustani, S., and Destexhe, A. (2007). Simulation of networks of spiking neurons: A review of tools and strategies. Journal of Computational Neuroscience, 23(3): 349–398.

Fidjeland, A. and Shanahan, M. (2010). Accelerated simulation of spiking neural networks using GPUs. pages 1–8.

Goodman, D. and Brette, R. (2008). Brian: a simulator for spiking neural networks in python. Frontiers in neuroinformatics, 2:5.

Goodman, D. F. M. (2010). Code Generation: A Strategy for Neural Network Simulators. Neuroinformatics, 8(3): 183–196.

Goodman, D. F. M. and Brette, R. (2009). The Brian simulator. Frontiers in Neuroscience, 3(2): 192–197.

Hoang, R. V., Tanna, D., Jayet Bray, L. C., Dascalu, S. M., and Harris, F. C. (2013). A novel CPU/GPU simulation environment for large-scale biologically realistic neural modeling. Frontiers in Neuroinformatics.

Knight, J., Yavuz, E., Turner, J., and Nowotny, T. (2018). GeNN (version 3.2), https://doi.org/10.5281/zenod0.593735.

Mutch, J., Knoblich, U., and Poggio, T. (2010). CNS: a GPU-based framework for simulating cortically-organized networks. Computer Science and Artificial Intelligence Laboratory Technical Report.

Nageswaran, J. M., Dutt, N., Krichmar, J. L., Nicolau, A., and Veidenbaum, A. V. (2009). A configurable simulation environment for the efficient simulation of large-scale spiking neural networks on graphics processors. Neural Networks, 22(5–6):791–800.

Nowotny, T., Huerta, R., Abarbanel, H. D. I., and Rabinovich, M. I. (2005). Self-organization in the olfactory system: Rapid odor recognition in insects. Biol Cybern, 93: 436–446.

NVIDIA^®^ Corporation (2006-2018). CUDA^TM^, https://developer.nvidia.com/cuda-zone.

Oh, K.-S. and Jung, K. (2004). GPU implementation of neural networks. Pattern Recognition, 37(6): 1311–1314.

Rolfes, T. (2004). Neural networks on programmable graphics hardware. Charles River Media, Boston, MA.

Stimberg, M., Goodman, D. F. M., Benichoux, V., and Brette, R. (2014). Equation-oriented specification of neural models for simulations. Frontiers in Neuroinformatics, 8:6.

Stimberg, M., Goodman, D. F. M., and Brette, R. (2018a). Brian 2 (version 2.2), https://doi.org/10.5281/zenod0.1459786.

Stimberg, M., Nowotny, T., and Goodman, D. F. M. (2018b). Brian2GeNN (version 1.2), https://doi.org/10.5281/zenod0.1464116.

Traub, R. D. and Miles, R. (1991). Neural Networks of the Hippocampus. Cambridge University Press, New York.

Vitay, J., Dinkelbach, H. Ü., and Hamker, F. H. (2015). ANNarchy: a code generation approach to neural simulations on parallel hardware. Frontiers in Neuroinformatics.

Yavuz, E., Turner, J., and Nowotny, T. (2016). GeNN: A code generation framework for accelerated brain simulations. Sci. Rep., 6:18854.

